# White-box Deep Neural Network Prediction of Genome-Wide Transcriptome Signatures

**DOI:** 10.1101/2021.02.11.430730

**Authors:** Rasmus Magnusson, Jesper N. Tegnér, Mika Gustafsson

## Abstract

Prediction algorithms for protein or gene structures, including transcription factor binding from sequence information, have been transformative in understanding gene regulation. Here we ask whether human transcriptomic profiles can be predicted solely from the expression of transcription factors (TFs). To this end, we explore whether a neural network (NN) could predict the transcriptome from TFs. Using at least one hidden layer, we find that the expression of 1,600 TFs can explain >95% of variance in 25,000 genes. Using the light-up technique to inspect the trained NN, we find an overrepresentation of known TF-gene regulations. Furthermore, the learned prediction network has a hierarchical organization. A smaller set of around 125 core TFs could explain close to 80% of the variance. Interestingly, reducing the number of TFs below 500 induces a rapid decline in prediction performance. Next, we evaluated the prediction model using transcriptional data from 22 human diseases. The TFs were sufficient to predict the target genes’ dysregulation (rho=0.61, P < 10^−216^). By inspecting the model, key causative TFs could be extracted for subsequent validation using disease-associated genetic variants. In conclusion, we demonstrate the construction of an interpretable neural network predictor. Analysis of the predictors revealed key TFs that were inducing transcriptional changes during disease.

## Introduction

To bridge the gap between genome sequences and phenotypes, it is essential to characterize intermediate levels, such as cells, tissues, and organs, using a suite of molecular technologies. For example, genetic variants associated with diseases exert their effects through the modulation of these intermediate levels. The transcribed mRNA expression is one of the most accessible and important windows into not only the regulatory machinery of the cell but also of changes in tissues and organs. Therefore, the analysis of mRNA expression is of crucial importance for the study of diseases [1]. Specifically, the elucidation of gene regulatory mechanisms is central since gene regulatory networks maintain cellular identity and mediate interactions between genetic variants and the environment of humans.

To reverse-engineer gene regulatory mechanisms, large amounts of RNA expression data have been generated from experimental model systems, including cell-lines and tissues from humans under different conditions [2]. Bioinformatics analysis has been instrumental in clustering genes to make sense of such data, thereby providing a hypothesis of genes involved in diseases [3]. Enrichment and pathway analysis increase the resolution by suggesting groups of genes or specific pathways associated with the observed changes in gene expression. Since transcription factors (TFs) are critical for the regulatory control of genes, a massive body of bioinformatics tools targets TF binding sites’ predictions, suggesting key drivers behind pathways, groups of genes, or clusters [4]. While useful for descriptive purposes, such as associating such differentially expressed genes to many diseases, it is challenging to gain functional and mechanistic insight into the regulatory machinery from such lists.

To advance beyond lists, clusters, and enrichment analysis, a complementary strategy, referred to as network science, instead targets the study of interactions between molecular entities, genotypes, and phenotypes [5], [6]. For example, gene regulation effectively acts via a network of interacting genes [7]. Notably, genes that interact with dysregulated genes without being differentially expressed themselves are often overlooked in differential expression studies [8]. Consequently, these networks are challenging to extract from data [9]. The wisdom of the crowd strategy has turned out to be useful while not satisfactory [10]. The limited progress originates from the fact that the inverse problem of inferring interactions from observations is statistically under-constrained. Moreover, these approaches have all struggled with the complex and nonlinear dynamics that shape gene regulation, containing several saturation effects and abundant negative and positive feed-backs. These non-linearities impede most of the available correlation-based methods used to study gene expression [11].

Recent progress in machine learning has fueled interest in whether such methods could facilitate the discovery and analysis of biological networks [12], [13]. Pioneering applications of deep neural networks (DNNs) in genomics include prediction of TF binding sites [14]), and the effects of non-coding genetic variants [15]. At the core of these techniques is the ability to capture non-linear relationships. The use of DNNs requires substantial amounts of data, which is now feasible due to the collection of massive amounts of genetic and RNA-Seq data into easily accessible databases [2]. Beyond detection of features, recent use of DNN, such as deep autoencoders applied to transcription data, compresses gene expression data into a latent space. The original data can then be reconstructed from the latent space representation [16]. DNNs have also been applied to understand the regulation of mRNA expression. Deep convolutional neural networks could predict 59% of human RNA abundance variation from the genomic sequence alone [17]. While being a first important step towards predicting mRNA levels, the regulatory transcription factors were not separated from the remaining transcriptome, making a biological interpretation and translation to diseases challenging.

Here we develop a methodology that goes beyond producing lists of differentially expressed genes, but not so far as the yet intractable reconstruction of a complete gene regulatory network. Instead, we target the regulation exerted by the transcription factors and ask whether training a DNN on gene expression data could learn a predictive TF network. The expression of all other genes could be explained by the combinatorial control induced by the TFs. Importantly, we constraint the training such that the resulting predictive model is interpretable, here referred to as a white-box model in contrast to a black-box machine learning model. We find that such models can indeed predict the expression of genes based on TFs and that the predicted relationships between TFs and their target genes largely overlap with known TF bindings. We apply and evaluate this predictive model using human disease transcriptomes, thus opening the door for a mechanistic and interpretable machine learning analysis of the human gene regulation system.

## Results

First, we explore the influence of different neural network architectures, such as the number of layers and hidden nodes on predictive performance. Next, we ask to what extent we can inspect the trained network and disentangle the different predictors’ contributions (TFs). Using a light-up network analysis technique, we identify a core set of TFs, including some key master regulators. In the final section of the results, we ask whether the prediction model could analyze disease derived transcriptional data.

### Accurate and robust prediction of the expression level of target genes using deep neural transcription factor networks

To optimize the performance of the applied Deep Neural Network (DNN), we compared 15 DNN architectures consisting of 1-3 intermediate layers, each with a depth of 50-1,000 hidden nodes. As a reference, we used a shallow NN without any intermediate layer (Fig. 1). The rationale is to identify the most compact architecture, measured by depth and width, capable of predicting most of the target gene expression with sufficient accuracy. Here, TFs were used exclusively as input and non-TFs as output target genes. We extracted TFs using the compendium provided by Lambert et al. 2018 [18], which lists TFs based on several sources, including popular TF databases such as TRANSFAC, JASPAR, and HT-SELEX. We trained models using the ARCHS4 database, using more than 100,000 randomly drawn RNA-Seq samples to train and evaluate the models [2]. The performance was evaluated using the gene-specific coefficient of determination (R^2^) on test data. We observed median R^2^ values between 0.88 and 0.97 depending on the model (Fig. 2a-b). Similarly, the mean R^2^ values of all DNNs were in the range of 0.89-0.93, while the shallow model had a notably worse performance (R^2^=0.75).

**Figure 1:**
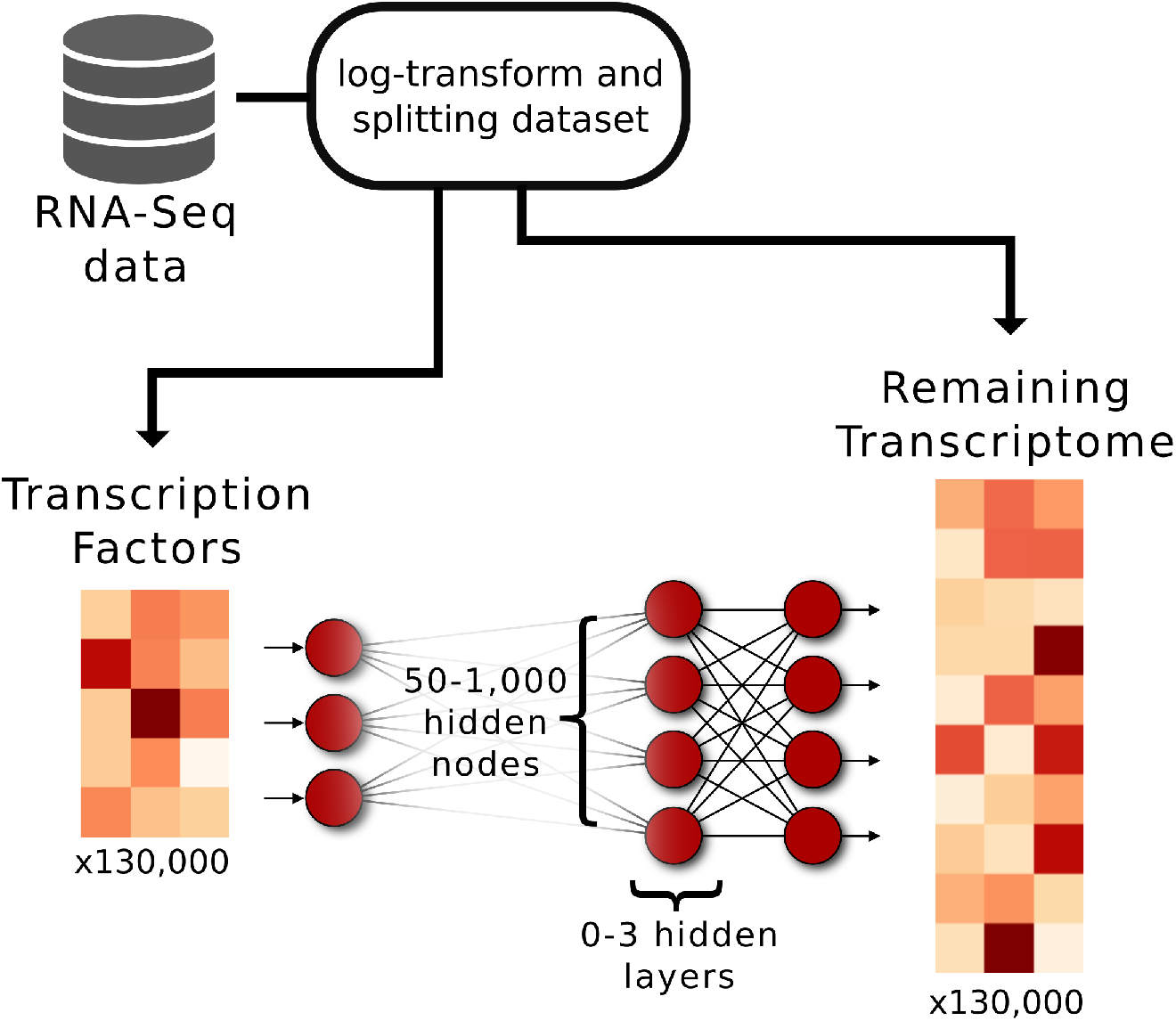
Design of the artificial deep neural networks. The networks were made to predict the expression of 25,861 genes based on 1,625 gene regulators, i.e. transcription factors (TFs). We used more than 100,000 randomly drawn RNA-Seq samples from the ARCHS4 database to train the models. Moreover, we designed 15 DNNs of 1-3 hidden layers, and one shallow neural network without any hidden layers. For the DNNs, each hidden layer consisted of either 50, 100, 250, 500, or 1,000 hidden nodes using the exponential linear unit, ELU as activation function.

**Figure 2:**
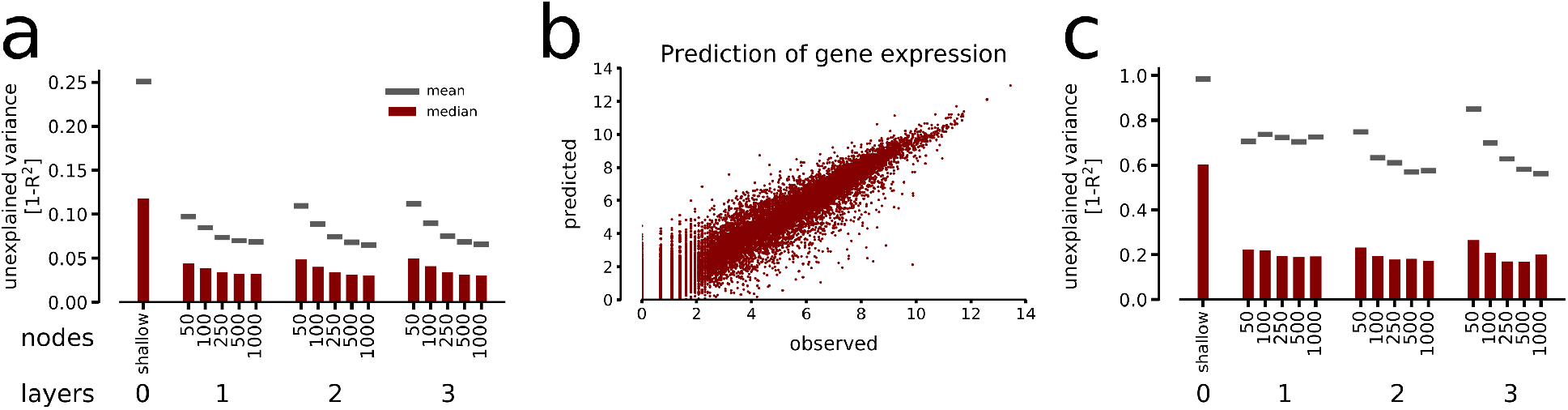
Gene expression prediction performance. a) We applied the DNNs and the shallow model to previously unseen gene expression profiles randomly selected from the ARCHS4 database, and calculated the coefficient of determination, R^2^, for each gene. Shown are the median (red bar) and mean (gray line) of the R^2^ values for each model. b) The typical prediction of an experiment is shown for the DNN with two hidden layers and 250 hidden nodes in each layer. c) We applied the DNNs to predict gene expression values from 934 human tumor samples from the Cancer Cell Line Encyclopedia. Shown are the abilities to predict this data set for the respective models, following the same layout as in a).

To test the trained models’ generalizability, we applied them to predict expression from the independent Cancer Cell Line Encyclopedia resource [19], which contains mRNA profiles of 934 human tumor cell lines. In this out-of-sample prediction, the DNNs showed a high median R^2^ between 0.74 and 0.83, whereas the shallow model only reached a median R^2^ = 0.40. Interestingly, the mean R^2^ values were lower than the medians for all models (0.15-0.44), resulting from the strong perturbation of the TF regulatory circuits that may occur in cancer. Nevertheless, the non-linear DNNs could capture such perturbations almost three-fold better than the shallow NN, showing their potential usefulness even in cancer medicine. With these results, we concluded that the DNNs could faithfully predict the majority of the human transcriptome given the expression levels of ~1,600 TFs, both in healthy and disease-affected states.

### Enrichment of validated TF-target associations within the prediction networks were revealed by the node light-up technique

Here we ask whether the trained networks are interpretable, that is, whether the learned TF target associations are biologically relevant or not. From a biological standpoint, one could argue for a direct comparison with known physical DNA bindings between TFs and their target genes. However, due to the non-linear dependencies embedded in a DNN, such investigations are not trivial [20]. We, therefore, reasoned that in the predicted network, TF expression perturbations would propagate most effectively to relevant target genes. Such an analysis captures the effective gene dependencies, linear or non-linear. To this end, we used a node light-up analysis. In other words, the numerical value of each input node, corresponding to a unique TF, was independently perturbed to either half or double of the mean gene expression. The other TF expression values were clamped to their average values, while the responses on the output layer were ranked by the response to such a TF change (Fig. 3a). To validate the DNNs, we compared the light-up values to previously known TF-target bindings. Specifically, we tested whether the light-up responses between such TF-target pairs were significantly higher than the pairs not annotated as interactions.

**Figure 3:**
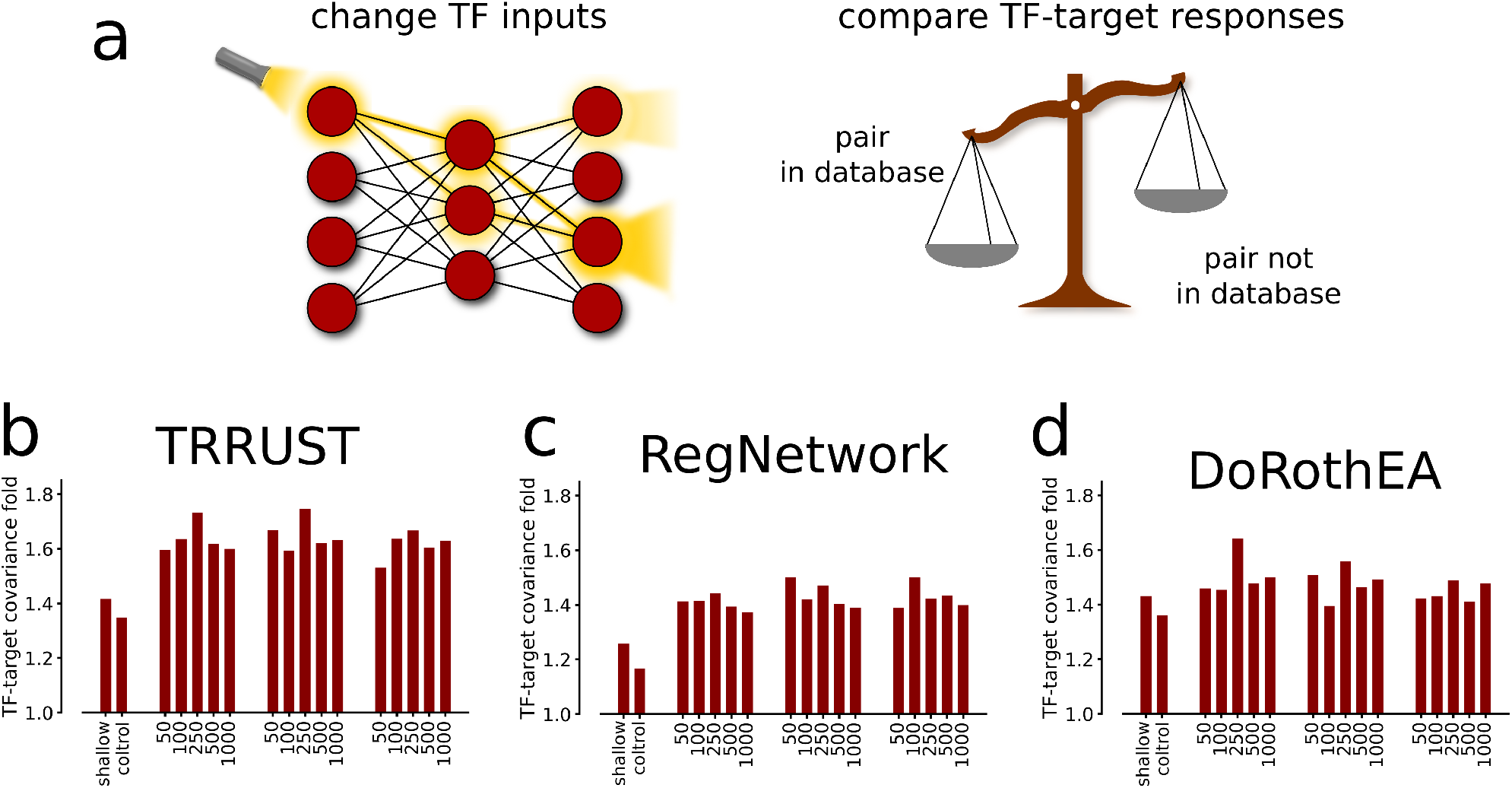
Node light-up reveals known TF-target associations. a) By applying a light-up analysis, i.e. by changing the input values of each TF independently and subsequently observing the changes on the output layer, we were able to estimate how the TF-to-target mappings corresponded to TF bindings known from literature. We defined the TF-target covariance fold as the median value of the light-ups for the TF-target regulations found in a database divided by the respective backgrounds. b) The light-up enrichments for the TF-target associations found in the TRRUST database are shown. Note that the expected value representing no biological relevance between TF-target mapping is 1 and that light-up values are compared to absolute Spearman rank correlation values, labeled ‘control’. We performed the same analysis with RegNetwork (c), and the interactions annotated with top confidence in DoRothEA (d).

We used three sources of experimentally validated TF-target interactions. Specifically, the TF-specific DoRothEA database (*n_TFs_* = 94, *n_targets_* = 2,240, *n_edges_* = 4,498) and TRRUST database including a better TF coverage (*n_TFs_* = 481, *n_targets_* = 1,961, *n_edges_ =* 6,576), and the RegNetwork database, which includes many more interactions of somewhat less confidence (n_TFs_ = 645, *n_targets_* = 14,696, *n_edges_* = 99,488). We also noted a limited overlap between these three databases. For example, 927 interactions were shared between the TRRUST and DoRothEA sets, 1,262 between DoRothEA and RegNetwork, and 751 between RegNetwork and TRRUST. Strikingly, the light-up responses between known TF-target associations were significantly higher (26-75%) for all databases and DNNs (binomial P – 10^−102^ in all cases). While the results using the three databases did not point to a single best model architecture, we observe the highest overall performance for the DNN with two hidden layers using 250 nodes in each layer. The highest enrichment rank was obtained from TRRUST, while DoRothEA yeilded the second-highest, and RegNetwork the lowest rank.

Since TFs can act as both inhibitors and initiators of transcription, we asked the extent to which the DNN light-up analyses also captured the directionality and sign of the TF-target interactions. Therefore, we compared the light-ups with known interactions, directions, and signs of TRRUST and DoRothEA. Of note, RegNetwork does not contain any annotation of interaction signs. Again, all tested DNNs showed significant overlaps with the interaction signs of both databases, with accuracies ranging between [0.62-0.67] (P ∈ [10^−120^,10^−58^]) for DoRothEA, and [0.56-0.58] (P ∈ [10^−22^, 10^−11^]) for TRRUST. We defined accuracy as the percentage of correctly estimated sign of interaction. Again, the shallow model performed significantly worse and could only predict interaction signs compared to DoRothEA (accuracy= 0.57, P< 10^−20^).

In general, we find that the number of hidden layers or units for the DNNs has only a limited impact on the performance. All DNN models showed comparable performance across evaluative analysis, expression predictions of experiments from the ARCHS4 database, the cancer expression predictions, and the light-up comparison to TF-target databases. In practice, 250 hidden nodes in two hidden layers appeared to be optimal in model size. Notably, the shallow NN never reached a satisfactory performance. Thus, the largest gain in explanatory power and overlap with existing databases came in our hands from adding at least one intermediate layer, thereby allowing for non-linear transformations. While these non-linearities turned out to be essential for the performance, they did not prevent us from inspecting the predictive network and extract relevant and validated biological knowledge using the light-up techniques.

### Algorithmic extraction of a core set of validated master regulator TFs from the DNN

We next searched for a minimal subset of key TFs required for predicting the target gene expressions, commonly referred to as master regulators. For this purpose, we implemented a backward-selection algorithm to stepwise remove TFs from the input layer based on their explanatory power (Methods). We observed highly consistent orders of TF-removal between the models, again suggesting that the TF-target relationship is robust to different DNN architectures. Indeed, the median correlation between different models, estimated from which step the TFs were removed, was 0.71 (geometric mean P < 10^−364^). This suggests a high consistency between the independently trained models (Supplemental Material S1). This observation, taken together with the light-up analysis outcome, suggests robustness of our results in that specifics of a DNN have only a minor influence on the results. Therefore, in the analysis’ remaining parts, we used the model with two hidden layers and 250 hidden nodes.

As expected, the explanatory power gradually decreases when removing predictors, here TFs (Fig. 4a). However, the shape of this loss of explanatory power exhibited two distinguishing features. Firstly, the ability to predict mRNA expression remained relatively unperturbed even as most TFs were removed from the set. Indeed, for n*_TFs_* = 125, the mean R^2^=0.79 compared to that of the full model of 1,625 TFs, which measured 0.93. Secondly, there appeared to be a stratification of TFs based on how important they are to explain the system, leading to a larger reduction in explanatory power towards the procedures’ late stage.

**Figure 4:**
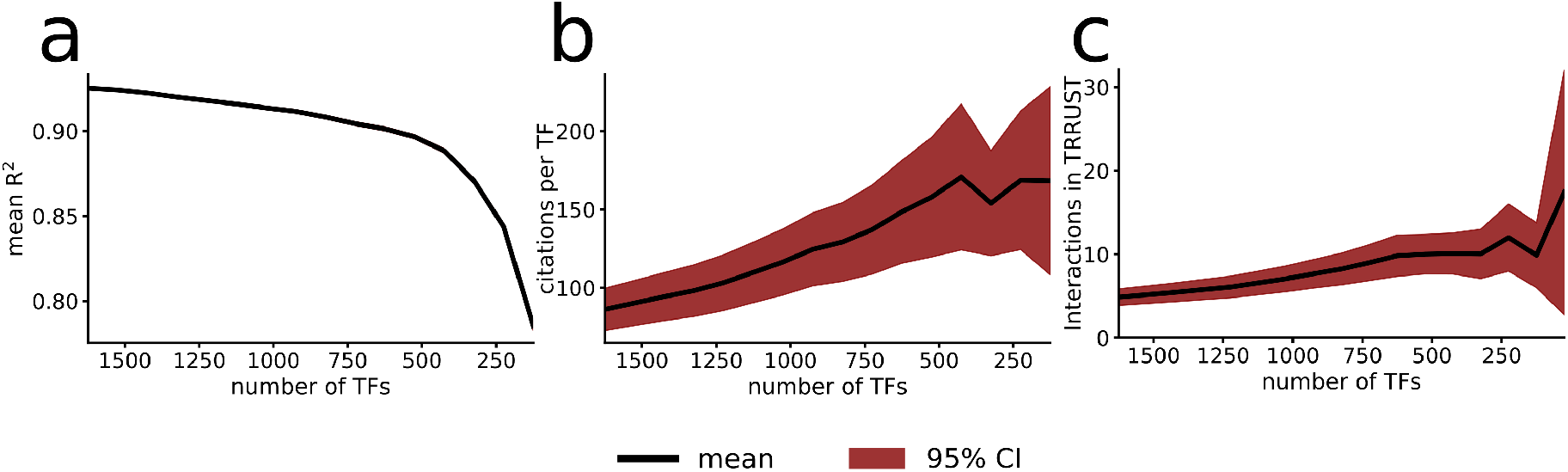
Functional Hierarchies in Model Input. a) We found the explanatory power on test data to decrease as the number of TFs were removed from the model. Particularly, we found around 500 TFs to carry a greater predictive power, as the explanatory power after that point more rapidly decreased. b) We measured the median number of scientific studies at each iteration of our backward-selection algorithm, and found that as the number of input TFs decreased, the more studied were those that were left. c) The TFs that were identified as more important also had more recorded interactions in the TRRUST database.

To test whether our predicted core set of TFs could correspond to known master regulator TFs, we estimated how often or much they have been studied in the literature domain. First, we analyzed the average number of scientific publications per input TF, as annotated by PubMed [21], and found that the top predicted TFs were significantly more frequently studied (Fig. 4b). For example, the median number of publications was 51 for the top 125 TFs, compared to 24 for all TFs (Mann-Whitney U test P < 8.2 * 10^−9^). Second, we analyzed the light-up conformity to the TRRUST database, but this time for each step in the backward-selection algorithm. We found that the full model performed equally well or better at next-to-all steps in the backward-selection algorithm. This observation indicates that the model predominantly discovered known TF-target interactions regardless of model input size. Third, the number of annotated interactions per TF was significantly larger for the top-ranked TFs in the literature-based database TRRUST (9.9 for top 125 compared to 4.9, P < 2.4 * 10^−6^), suggesting that the top explanatory TFs were associated with more well-known regulatory bindings. We next analyzed the functional role of the removed TFs and performed enrichment analysis of their predicted downstream targets from our light-up procedure (Methods). For each set of 100 removed TFs using the backward-selection procedure, we associated their top 500 target genes, which were then subjected to a KEGG pathway [22] enrichment analysis. In brief, we found that the top 100 TFs (i.e., the last removed TFs) were mostly enriched for cellular metabolism pathways, followed by immune-pathways for the TFs with rank 100-200 (Supplemental Table S2). Thus, despite having abundant input variables, the DNN discovered known core TFs associated with central and well-annotated pathways.

### Latent DNN Space Shows enrichment of Functionally Related and Disease-Associated Genes

In addition to assessing the biological relevance of the specific interactions discovered by the predictive DNN, one may ask whether the predictor could be informative in a disease context. It has been demonstrated that compressing mRNA expression data in deep autoencoders can provide low-dimensional representations exposing complex characteristics of the input data domain. Importantly, from such a representation, one can readily extract sets of functionally related genes, known as modules, which in turn can be used to study diseases [16]. To further increase the resolution beyond the analysis of modules, here we aimed to use the TF-target gene interactions within the two 250 variables measuring intermediate latent layers.

Specifically, we tested whether the genes associated with these hidden variables shared cellular functionality. For this purpose, we again used a light-up response to associate genes to each hidden node independently. We annotated the top 500 responding genes to each node light-up by this procedure and performed a Reactome pathway enrichment analysis for each node-set. This procedure identified 175 unique Reactome pathways that significantly overlapped with at least one hidden node (using a Bonferroni correction of 0.05). In detail, 102 hidden nodes of the first and 162 of the second layer were associated with at least one Reactome pathway. These results suggested at least a subset of the hidden nodes represented different cellular pathways. We evaluated the putative agglomeration of disease-related genes in the hidden-layer light-up responses. To this end, using each hidden node, we performed a genome-wide association study (GWAS) enrichment analysis between annotations in the NHGRI-EBI GWAS catalog [23] and the 500 genes with the highest light-up associations. We found 37 of the 153 diseases associated with one or several nodes, with 63 nodes in the first and 62 nodes in the second layer having at least one association. Second, we cross-checked these results by comparing the light-ups with the DisGeNET database [24], which contains broader profiles of gene-disease associations. We tested against genes grouped in 26 disease-type categories and found 22 of these categories enriched with at least one hidden node. At least one significant overlap with a disease category was detected in 107 nodes in the first and 138 nodes in the second layer. Thus, disease-genes appeared to co-occur in hidden-node light-ups in the DNN. This result suggests that the remaining genes found in such disease modules could be relevant in analysing and interpreting disease-related biomarkers and mechanisms.

### DNN Analysis gives Insights to Human Disease Mechanisms involved in Gene Dysregulation

We finally aimed to test the clinical relevance of the DNN by using it to study disease-related changes in the expression of target genes, given the corresponding modifications of TF levels. To this end, we applied the DNN to independent RNA-Seq data from the Expression Atlas [25] using differential expression patterns for diseases from 27 different studies, which together contained 69 expression fold profiles. We tested if the DNN could predict the differential expression of target genes given the fold profiles of the TFs. Prediction quality was measured as the correlation between the predicted and observed fold ranking of significantly differentially expressed target genes for each disease. In other words, we set the TFs to their reference expression levels and applied the fold changes of each respective disease. We next calculated the correlation between the observed and predicted fold changes at the output layer. Notably, we observed highly significant correlations between these predictions and observations, with a median Spearman rank correlation of 0.61 (median P < 10^−216^). This result established that disease mechanisms of dysregulation from TFs could be faithfully propagated to the target level. Yet, as a correlation does not imply causation, we assessed whether the DNN could be used to disentangle which TFs drive the target dysregulation. Replacing dysregulated with reference expression values independently for each TF and evaluating the impact on the output layer enabled us to rank the predicted TF impact as a change in correlation between predicted and observed target dysregulation (Fig 5). Next, we matched these rankings to known genetic variants from genome-wide association studies (GWAS) in the 22 applicable cases and measured the area under the receiver operating characteristics curve (AUROC). We found the TF rankings in 10 out of 22 diseases to significantly match the TF GWAS annotation (binomial test P < 3.6 * 10^−8^).

**Figure 5:**
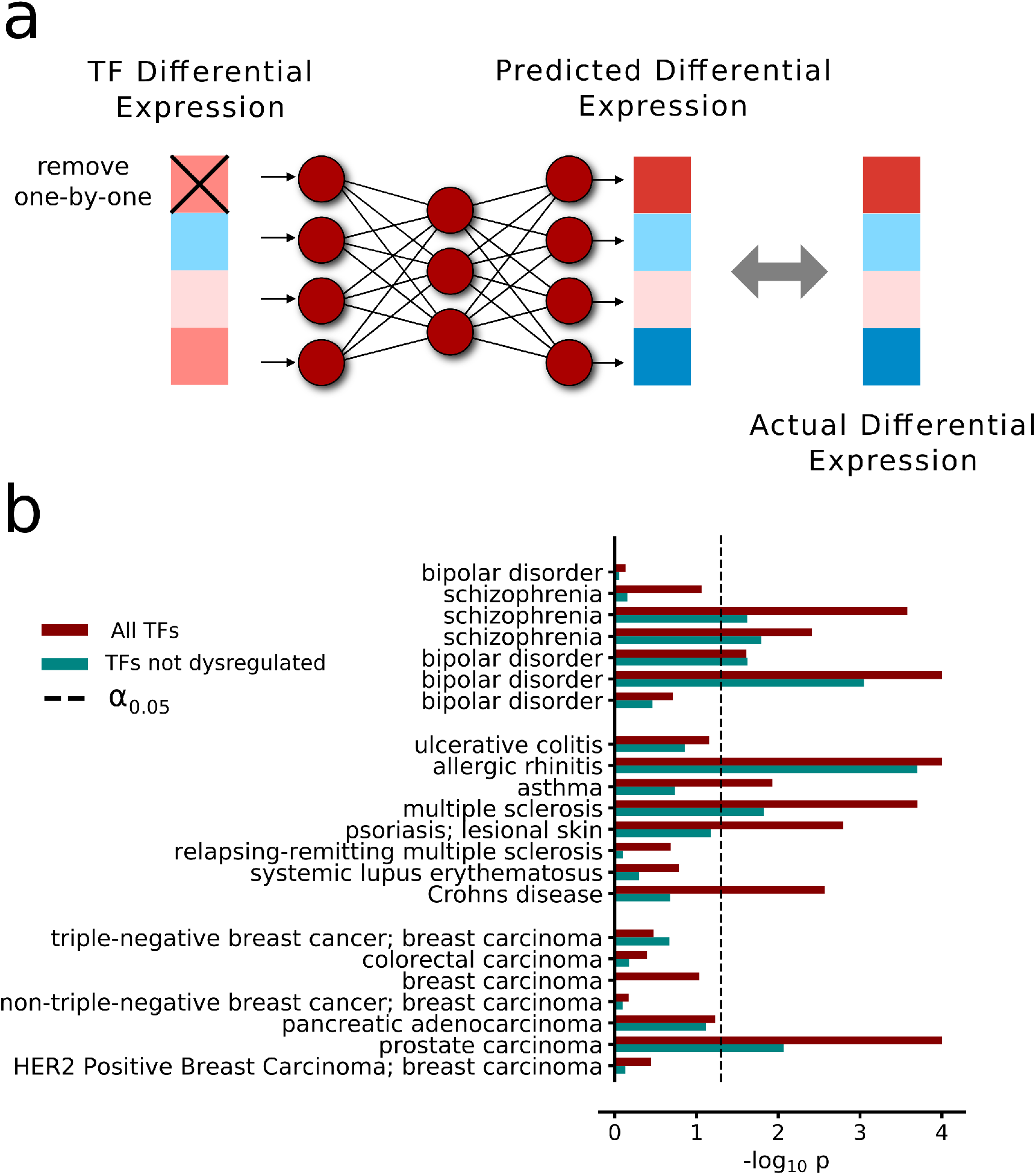
Predicting gene dysregulations in disease using a DNN. a) We tested if the DNN with two hidden layers of 250 hidden nodes each could be used to predict causative changes in disease states. We did this by analyzing gene expression changes from known diseases, as available in the Expression Atlas repository. By applying the disease changes to the transcription factor input layer, we could observe how these changes projected down to the target genes. Next, we removed the disease-fold changes of each TF independently and observed the changes in correlation between predicted and observed dysregulation of the target genes. Thus, we could rank the TFs on predicted causative disease-changes on the target genes. b) TF rankings significantly overlapped with GWAS in 10 of 22 diseases. (–*log_10_* P-values shown as red bars). The test was repeated with significantly differentially expressed TFs removed from the set, leaving 7 TF rankings to overlap with GWAS. (–*log_10_* P-values shown as teal bars).

Finally, we asked if TFs had to be differentially expressed to carry predictive power over disease-affected target gene regulation. This question is of particular importance since causative disease-related changes are not necessarily manifested through dysregulations that are large enough to be detected in multiple-testing corrected statistical tests of expression changes. Notably, we found our approach of causative predictions on a TF level to also give significant results when only applied to TFs that themselves were not significantly dysregulated, with seven diseases remaining significant (Fig 5, P < 6.8 * 10^−5^). This suggests that our approach is generally applicable to find disease-causative elements at the TF-level, beyond what a more conventional RNA-Seq analysis of gene expression detects. In summary, the TF-to-target DNN allowed a simple representation to describe the observed differential expression between patients of many different complex diseases and control in terms of TFs, which strikingly also correctly associated a truly enriched fraction of disease-associated TFs to several diseases.

## Discussion

We have presented a biologically interpretable, general machine learning method for predicting transcriptomic signatures, including disease signatures. Our learned models predict the expression of genes from the expression of transcription factors (TFs). The predicted relationships between TFs and their target genes largely overlap with known TF bindings. Hence, our DNN method goes beyond classic descriptive bioinformatic techniques such as clustering and enrichment analysis. Importantly, we do not address the yet intractable problem of complete deconvolution of the entire cellular interactome. Instead, our method does extract a core TF component out of such complex regulatory architecture. Therefore, the presented methodology sets the stage for the first step towards mechanistic and interpretable data-driven machine learning analysis of the human gene regulation system.

Early success of machine learning techniques targeted fundamental open bioinformatics problems such as predicting binding sites of TFs [14] or functional prediction of non-coding RNA in a disease context [15]). The recent extension has focused on questions such as if one data-type can be predicted from another data-type. Prediction of gene expression from the DNA sequence or prediction of the 3D genomic structure from open chromatin data are just two recent examples. Tasaki et al. 2020 [26] used deep convolutional neural networks to predict differential expression based on mRNA features and the promoter regions of the genes. Yet, here as in the other examples, these impressive advances are difficult to translate into biological knowledge. As in different machine learning areas, these systems are useful predictors but act in practice as black-box systems. Related to this is that even if functionally relevant representations can be identified, they can readily suffer from study-biases of identified prevalent genes. Remaining core challenges in using machine learning techniques include interpretability of the model, large data-requirement, and how to learn biologically meaningful representations within the machine learning model. A black-box model does not lend itself to interpretable and meaningful representations which may also make the model more susceptible for adversarial attacks [27], [28]. Recently, it has become increasingly clear that deep neural networks (DNNs) have the potential to identify biologically meaningful molecular representations directly from data [16], [29] and to revolutionize medicine [30], [31].

Our main contribution is to design a constrained machine learning approach such that the predictor is interpretable from a biological standpoint. Using two hidden layers with 250 hidden nodes each was sufficient to capture well-known TF-target pairs. These results suggest that the learned representation has significant architectural overlap compared to a proper cellular control system, which is essential for a good generalizability of DNNs. By developing a back-tracking algorithm, we could uncover a minimal core set of 125 TFs sufficient to account for ~80% of the transcriptomics signature. Interestingly, these TFs represent the TFs being studied most frequently, which makes further studies linking prior structural information from other data-types a logical next step to increase the generalizability even further.

Our focus on TFs originates from the observation that TFs have been at the forefront in analyzing cellular reprogramming and converting cell-types [32], [33]. Our findings suggest that the presented DNN methodology could be useful as an interpretable predictive method for pinpointing key candidate TFs for cellular reprogramming using large transcriptomics data [34]. One of the most common approaches to analyze disease mechanisms is the study of gene expression changes between healthy and disease-affected individuals. Traditionally, a suite of bioinformatics descriptive mining techniques is applied to extract putative candidates of interest [3]. Nevertheless, it has proven challenging to pinpoint molecular mechanisms with high precision using such data-mining approaches. Consider a scenario where the impact of a perturbation to the gene regulatory system is of interest. Such cases are a common end-goal of analyzing transcriptomics and pivotal to understand mechanisms such as drug perturbations or impacts of cancer [35]. Using our light-up analyses could help predict specific regulatory interactions and their effect on the transcriptome. This could be useful both for cellular reprogramming experiments as well as in the analysis of diseases. In contrast, a machine learning model using only gene sequences would not readily consider such changes.

Lastly, since our predictor’s architecture has a biological interpretation, it could be used as a first approximation – like a blueprint – of the regulatory networks controlling the cell-identity and filtering effects of genetic variants. This problem has been at the forefront in systems biology since the sequencing of the human genome [10], [36]. Yet, despite two decades of brilliant work on reverse-engineering gene regulatory networks from data, it remains an open challenge [37]. The combinatorial complexity of such a network exceeds the amount and quality of available data given the current suite of models [10]. As network predictions vary between methods, new tools have been developed to control the abundance of false interactions [38], illustrating that the problem is still outstanding. Yet, if robust methods, such as our proposed DNN technique, could elucidate the TF part of such a network, we could potentially approach the problem in a step-wise manner. Hence, in summary, a TF-centric reverse-engineering technique could therefore be a stepping stone for renewed systems biology efforts in elucidating the cellar regulatory machinery at scale.

## Materials and Methods

### Data processing

We trained the models on gene expression data from ARCHS4, a database of >130,000 human RNA-Seq gene expression measurements from GEO and SRA. We separated the data into two sets, one of genes annotated as TFs and one containing the rest of the genes, which were assumed to be regulated downstream of the TFs. Moreover, we normalized the expression counts, annotated as x in Eq. (1), using the natural logarithm.

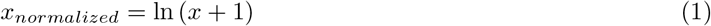

Moreover, we removed 1,200 gene expression profiles from the training set to use as validation of the model predictions. We next aimed to predict the expression levels of the target genes using the TF levels. We designed the DNNs to have the exponential linear unit (ELU) activation function on all nodes and across all layers. Furthermore, we set the Adam algorithm to minimize the mean squared error, as implemented in Keras.

### Backwards-selection algorithm

The algorithm operated according to the following three steps. First, each input node was independently set to zero, and the corresponding R2 values were calculated. Secondly, the bottom 100 TFs in terms of explanatory power were removed from the input layer of the DNN. Thirdly, we retrained the new and smaller model to explain the rest of the gene expressions. For each iteration, we tested the model on the same validation data from the ARCHS4-database as in the first validation experiment.

### Disease analyses

To analyse diseases, we used the Expression Atlas [25] to download all data according to the following criteria. I) The data was to come from a study carried out in human material. II) We only considered RNA-Seq experiments. III) The data sets had ‘disease’ listed as an experimental variable, and be of the ‘differential’ type. This query yielded data from a total of 27 studies, which together contained 69 expression fold profiles. Of these 69 comparisons, 56 were between a disease-affected and a healthy state, as opposed to between two disease-states, and we continued with these 56 studies. The 56 studies contain fold-changes between the healthy and disease affected states, and by adding these changes to the mean expression values from the ARCHS4 database we could predict the fold changes on the target gene level. Furthermore, we chose to only study the correlation between the genes that were differentially expressed

The predictions of TFs causative of disease were done by adding the fold changes to the TFs as described above, followed by removing the dysregulation of each TF independently. This resulted in 1,625 predicted changes in correlation between predicted and measured target expressions as compared to that from the full TF profile. We ranked the TFs on change in correlations and calculated the area under the receiver operating characteristics curve, i.e. the AUROC, for this ranking. We next Monte Carlo sampled 10,000 random TF permutations and estimated the P-value from the random AUROC distribution.

## Acknowledgements

This work was supported by the Swedish Research Council (grant 2019-04193(M.G.), the Swedish foundation for strategic research (grant SB16-0095(R.M., M.G.)), the Center for Industrial IT (CENIIT)(R.M., M.G.), the Systems Biology Research Centre at University of Skövde under grants from the Knowledge Foundation (grant 20200014) (R.M.), and the King Abdullah University of Science and Technology (KAUST) (J.N.T). Computational resources were granted by Swedish National Infrastructure for Computing (SNIC; SNIC 2020/5-177, and LiU-2019-25). The authors would like to thank Dr. Robert Lehman for his helpful suggestions to improve this manuscript.

## Code availability

The code used to generate the results, along with the DNN with 250 hidden nodes in two layers, can be found at https://github.com/rasma774/tf_target_white_box_dl.

## Notes

### Competing Interest Statement

The authors have declared no competing interest.

