## Supplementary material for "White-box Deep Neural Network Prediction of Genome-Wide Transcriptome Signatures": S1

### S1: Hierarchical structures of TF inputs are robust between models

For each of the models, we performed a backward-selection algorithm on the TFs (Methods) to rank the TFs according to importance when predicting target gene expression. These models were trained completely independently, and to analyse how robust TF-to-target interactions identified by the models were, i.e. how consistent the model approach was, we compared the similarities of removal order between the models. We measured similarity as correlation between the respective step at which each TF was removed, with one correlation between each model pair (Fig S1).

We found that the models that had a high compression, i.e. few hidden nodes in each layer, were less similar in the order TFs were removed. Nevertheless, all TF-ranking comparisons had highly significant Spearman correlations (highest P < 10e-118, rho=0.53).

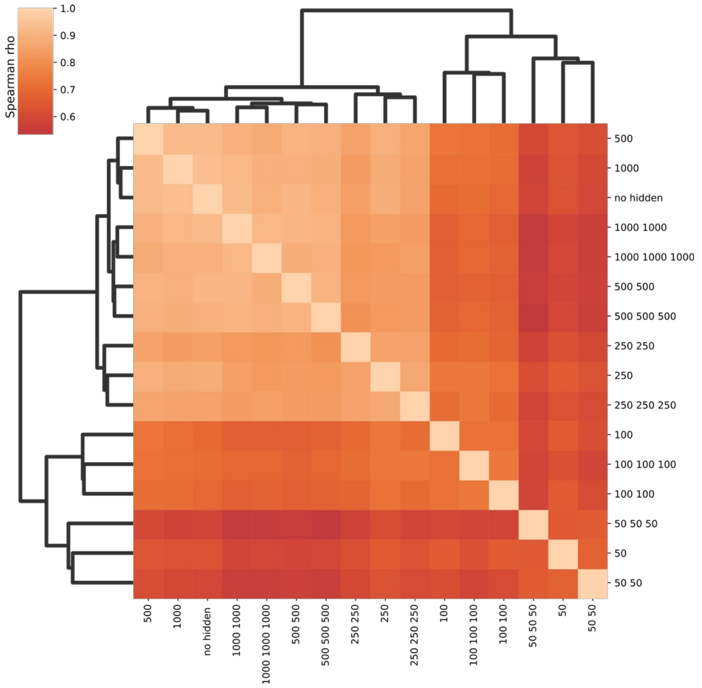


**Fig S1:** *Consistency of TF-removals between models. We found the iteration at which TF was removed to be highly consistent between models, indicating a robustness in how TFs are used to predict target genes.*
